# The Dot/Icm-translocated effector LegC4 potentiates cytokine-mediated restriction of *Legionella* within accidental hosts

**DOI:** 10.1101/493197

**Authors:** Tshegofatso Ngwaga, Alex J. Hydock, Sandhya Ganesan, Stephanie R. Shames

## Abstract

*Legionella pneumophila* is ubiquitous in freshwater environments where it replicates within unicellular protozoa. However, *L. pneumophila* is also an accidental human pathogen that can cause Legionnaires’ Disease in immunocompromised individuals by uncontrolled replication within alveolar macrophages. To replicate within eukaryotic phagocytes, *L. pneumophila* utilizes a Dot/Icm type IV secretion system to translocate a large arsenal of over 300 effector proteins directly into host cells. In mammals, translocated effectors contribute to innate immune restriction of *L. pneumophila*. We found previously that the effector LegC4 is important for *L. pneumophila* replication within a natural host protist but is deleterious to replication in a mouse model of Legionnaires’ Disease. In the present study, we used cultured mouse primary macrophages to investigate how LegC4 attenuates *L. pneumophila* replication. We found that LegC4 enhanced restriction of *L. pneumophila* replication within macrophages activated with tumor necrosis factor (TNF) or interferon (IFN)-γ. Specifically, TNF-mediated signaling was required for LegC4-mediated attenuation of *L. pneumophila* replication within macrophages. In addition, expression of *legC4* was sufficient to restrict *L. longbeachae* replication within TNF- or IFN-γ-activated macrophages. Thus, this study demonstrates that LegC4 contributes to *L. pneumophila* clearance from healthy hosts by potentiating cytokine-mediated host defense mechanisms.

**Importance:** *Legionella* are natural pathogens of protozoa and accidental pathogens of humans. Innate immunity in healthy individuals effectively controls *Legionella* infection due in part to rapid and robust production of pro-inflammatory cytokines resulting from detection of Dot/Icm-translocated substrates, including effectors. Here, we demonstrate that the effector LegC4 enhances pro-inflammatory host restriction of *Legionella* from macrophages. These data suggest that LegC4 may augment pro-inflammatory signaling or antimicrobial activity of macrophages, a function that has not previously been observed for another bacterial effector. Further insight into LegC4 function will likely reveal novel mechanisms to enhance immunity against pathogens.

## Introduction

*Legionella* are natural pathogens of unicellular protozoa and accidental pathogens of humans that can cause in a severe inflammatory pneumonia called Legionnaires’ Disease, which results from uncontrolled bacterial replication within alveolar macrophages. To replicate within eukaryotic phagocytes, *Legionella* subvert normal endocytic signaling by establishing a specialized compartment called the *<u>L</u>egionella* <u>c</u>ontaining <u>v</u>acuole (LCV). To form the LCV and replicate intracellularly, *Legionella* employ a Dot/Icm type IV secretion system (T4SS) to translocate virulence factors - called effector proteins - into host cells (1). Although >15 *Legionella* species are capable of causing human disease, the overwhelming majority is caused by *L. pneumophila* (2, 3). In healthy individuals, *L. pneumophila* infection is efficiently controlled and human-to-human transmission is incredibly rare (4). This is due to efficient detection and subsequent clearance of *L. pneumophila* by the mammalian innate immune system. Consequently, *L. pneumophila* is a well-established model pathogen used to characterize mechanisms of host defense against bacterial pathogens.

Innate immune detection of bacterial pathogens is facilitated by host pattern recognition receptors (PRRs) that recognize pathogen associated molecular patterns (PAMPs). Surface toll-like receptors (TLRs) are PRRs critical for host defense against *L. pneumophila*. The majority of TLRs signal through the adaptor MyD88 to activate pro-inflammatory gene expression. Mice lacking MyD88 are highly susceptible to *L. pneumophila* infection, which is mostly due to lack of interleukin (IL)-1 and TLR2-mediated signaling (5-7). Intracellular PRRs such as Nod1, Nod2, and inflammasomes also contribute to innate immune restriction of *L. pneumophila* in macrophages [reviewed in (8, 9)]. Leakage of PAMPs through the Dot/Icm pore amplifies cell-autonomous restriction of *L. pneumophila* within immune phagocytes. Specifically, recognition of *L. pneumophila* flagellin monomers (*flaA*) by the NAIP5/NLRC4 inflammasome is sufficient to restrict *L. pneumophila* replication within macrophages (10, 11). In addition, translocation of peptidoglycan, nucleic acids, and lipopolysaccharide into the host cell cytosol through the Dot/Icm pore also enhances restriction of *L. pneumophila*. Engagement of both extracellular and intracellular PRRs results in a robust pro-inflammatory response mediated by secretion of cytokines by infected and bystander immune phagocytes (5, 12-16). In particular, tumor necrosis factor (TNF) and interferon (IFN)-γ are critical for restriction of pulmonary *L. pneumophila* infection (17-21). TNF and IFN-γ both promote cell autonomous defense against *L. pneumophila* within macrophages and mediate bacterial killing by increasing phagolysosmal fusion (22, 23).

In addition to canonical PAMPs, translocated effectors can augment pro-inflammatory responses in *L. pneumophila*-infected macrophages. For example, effector-mediated inhibition of host protein translation results in increased expression of pro-inflammatory genes in macrophages (24-27). In addition, macrophage pro-inflammatory gene expression was decreased during infection with *Legionella* that possess a functional Dot/Icm T4SS but are unable to translocate a subset of effectors due to a mutation in the *icmS* effector chaperone gene (28). These studies elaborate the concept of effector-triggered immunity in animal cells (29) and provide further evidence for the contribution of effectors to innate immune restriction of *L. pneumophila*.

We recently demonstrated that the effector LegC4 attenuates *L. pneumophila* fitness in a mouse model of Legionnaires’ disease (30). Loss-of-function mutation of the *legC4* gene conferred a fitness advantage on *L. pneumophila* in the mouse lung as evidenced by increased pulmonary bacterial burden and the ability to outcompete the wild-type strain (30). However, the *legC4* mutation had no effect on *L. pneumophila* replication in primary bone-marrow derived macrophages (BMDMs) and impaired replication within a natural amoeba host, *Acanthamoeba castellanii* (30). Furthermore, expression of *legC4* from a plasmid further attenuated *L. pneumophila* fitness in the mouse lung compared to the wild-type strain. Thus, we hypothesized that LegC4 is deleterious to *L. pneumophila* in the presence of cell-mediated innate immunity.

The present study was designed to determine how LegC4 augments restriction of *L. pneumophila* in mammalian hosts. Expression of *legC4* from a plasmid was sufficient to attenuate *L. pneumophila* replication in BMDMs, which relied on TNF secretion and subsequent signaling. Moreover, a Δ*legC4* mutant exhibited increased replication in cytokine-activated BMDMs. Interestingly, expression of *legC4* was sufficient to attenuate *L. longbeachae* replication within TNF- and IFN-γ-activated BMDMs. These results suggest that LegC4 enhances macrophage cell autonomous defense against *Legionella* by potentiating cytokine-mediated restriction.

## Materials and Methods

### Bacterial strains, plasmids, primers, growth conditions

*Legionella pneumophila* Philadelphia-1 SRS43 (30), SRS43 *flaA*::Tn (30), Lp02 Δ*flaA* (10), Lp03 (31), and *Escherichia coli* strains were gifts from Dr. Craig Roy (Yale University). *L. longbeachae* NSW 150 was a gift from Dr. Hayley Newton (University of Melbourne). *Escherichia coli* strains used for cloning (Top10, Invitrogen) and *L. pneumophila* mutagenesis [DH5α-λ*pir*, (32)] were maintained in Luria-Bertani (LB) medium supplemented with 25 μg mL^-1^ chloramphenicol (pJB1806 and pSN85) or 50 μg mL^-1^ kanamycin (pSR47s). *Legionella* strains were cultured on supplemented charcoal N-(2-Acetamido)-2-aminoethanesulfonic acid (ACES)-buffered yeast extract (CYE) and grown at 37°C as described (33). *L. pneumophila* Lp02 strains were maintained on CYE supplemented with 100 μg mL^-1^ thymidine. Liquid cultures were grown at 37°C with aeration in supplemented ACES-buffered yeast extract (AYE) as described (33, 34). When necessary, media were supplemented with 10 μg mL^-1^ chloramphenicol (plasmid maintenance), 10 μg mL^-1^ kanamycin (allelic exchange) or 1 mM isopropyl-β-D-1-thiogalactopyranoside (IPTG).

Where indicated, recombinant mouse interferon-γ (rIFN-γ; Thermo Fisher Scientific), recombinant mouse tumor necrosis factor (rTNF; Gibco), rat α-mouse TNF antibody (α-TNF; R&D Systems), or normal rat IgG control (Rat IgG; R&D Systems) were used at a concentration of 50ng mL^-1^.

A complete list of oligonucleotide primers used in this study is shown in Table 1.

**Table 1.**
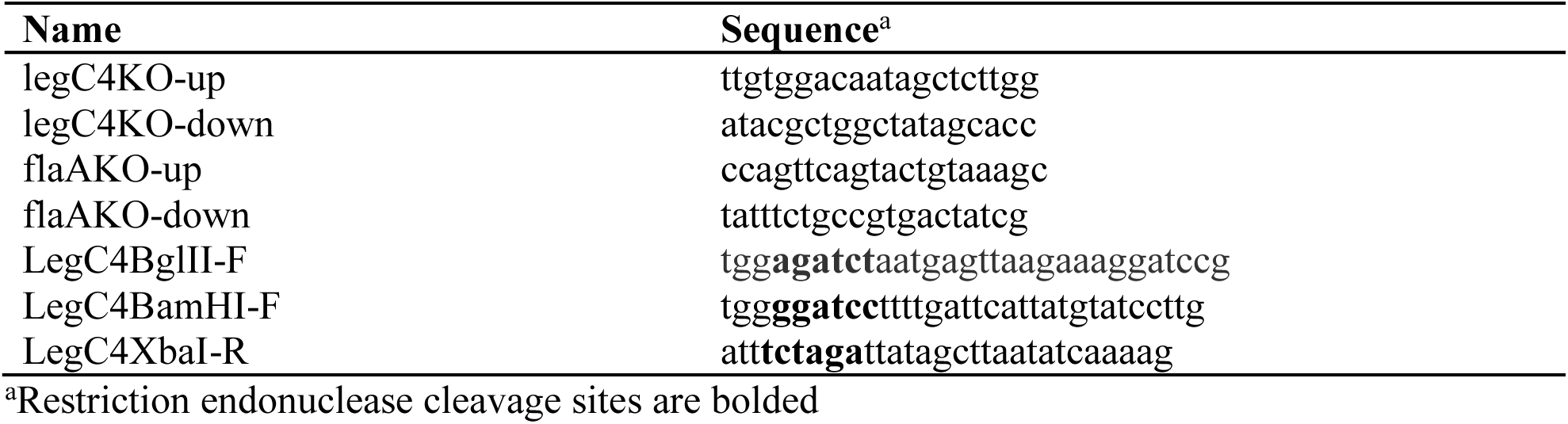
Oligonucleotide primers used in this study

### Molecular cloning, plasmid construction and generation of *Legionella* strains

In-frame deletions of *legC4* were generated by allelic exchange. Plasmids pSR47s:: Δ*legC4* and pSR47s::Δ*flaA*, gifts from Dr. Craig Roy, were conjugated into SRS43 or Lp02 followed by selection for *L. pneumophila* deletions as described (35, 36). Sucrose-resistant, kanamycin-sensitive colonies were screened by PCR using legC4KO-up/legC4KO-down and flaAKO-up/flaAKO-down primer pairs for Δ*legC4* and Δ*flaA* deletions, respectively.

To express *legC4* on a plasmid under its endogenous promoter, *legC4* plus the 300 base-pair upstream region was amplified with the LegC4BglII-F/LegC4XbaI-R primer pair and cloned as a BglII/XbaI fragment into BamHI/XbaI-digested pJB1806 (pJB) (37).

For IPTG-mediated expression of *legC4*, the *legC4* gene was amplified using the primer pair LegC4BamHI-F/LegC4XbaI-R and cloned as a BamHI/XbaI fragment into BamHI/XbaI- digested pSN85, a gift from Dr. Craig Roy (pEV; N-terminal 3XFLAG epitope tag fusion (38)). Sequence-confirmed pJB1806::p*legC4* (pJB::*plegC4*) and pSN85::*legC4* (p*legC4*) plasmids and empty vectors were transformed into *Legionella* strains as previous described (39). IPTG-induced expression of *3xFLAG-legC4* was confirmed by Western blot analysis (data not shown).

### Mice and BMDMs

C57BL/6 wild-type, *Myd88-/-*, and *Tnfr1-/-* breeding pairs were purchased from the Jackson Laboratories (Bar Harbor, Maine) and in-house colonies were maintained in specific pathogen-free conditions at Kansas State University. All experiments involving animals were approved by the Kansas State University Institutional Animal Care and Use Committee (protocol #4022) and performed in compliance with the Animal Welfare Act and NIH guidelines.

Bone marrow was harvested from mice as previous described (40). Bone marrow-derived macrophages were generated by differentiation in RPMI supplemented with 20% heat-inactivated fetal bovine serum (HI-FBS) (Gibco) and 15% L929 cell supernatant for 6 days prior to seeding for infection.

### Competitive index (CI) experiments in mice

Six-to ten-week old age and sex-matched C57BL/6 mice were infected for competitive index experiments as previously described (30). Mixed bacterial inoculums (1:1) were diluted and plated on selective medium (5 μg mL^-1^ for *flaA*::Tn and 10 μg mL^-1^ for plasmid selection). At 48 h p.i., mice were euthanized and whole lung tissue was harvested. Lung tissue was homogenized in 300 μL of sterile water using a Bullet Blender (Next Advance) as described (41) and dilutions were plated on selective medium as above. Colony forming units (CFUs) were enumerated and used to calculate CI values [(CFUcm^R^_48h_/CFUwt_48h_)/(CFUcm^R^_IN_/CFUwt_IN_).

### Quantification of *Legionella* replication within macrophages

Differentiated BMDMs were maintained in RPMI supplemented with 10% HI-FBS (Gibco) and 7.5% L929 cell supernatant. BMDMs were seeded 2.5 x 10^5^/well in 24-well plates one day prior to infection. BMDMs were infected with the indicated strains of *L. pneumophila* or *L. longbeachae* at a multiplicity of infection (MOI) of 1 in the presence or absence of 1 mM IPTG and/or recombinant cytokine as indicated. At 1 h p.i., cell monolayers were washed three times with PBS^-/-^ and fresh supplemented medium was added. Infections were allowed to proceed for up to 72 h or for 48 h, as indicated. To enumerate CFUs, BMDMs were lysed in sterile water for 8 mins followed by repeat pipetting. Lysates were diluted as appropriate and plated on CYE agar plates, which were then incubated at 37° for 4 days. For growth curve experiments, bacteria were enumerated after 1 h of infection and every 24 h thereafter for up to 72 h. To quantify fold replication, BMDMs were infected for 1 h and 48 h and fold replication was enumerated by normalization of the 48 h CFU counts to the 1 h CFU counts.

### Enzyme-linked immunosorbent assay (ELISA)

BMDMs were seeded in a 24-well plate at 2.5 x 10^5^/well 1 day prior to infection. The indicated Lp02 or SRS43 strains were used to infect the BMDMs (*n* = 3) at an MOI of 30 or 10, respectively. Infections with Lp02 strains were performed in the absence of exogenous thymidine to prevent bacterial replication. One-hour p.i., media were aspirated and cells were washed 3 times with PBS. Media were replaced and supernatants were collected after 5 h. Supernatants were either used fresh or stored at -20°C for up to 1 week followed by quantification of TNF using a Mouse TNF ELISA Kit (BioLegend) following manufacturer’s instructions.

### Western blot

To confirm production of 3XFLAG-LegC4 protein from *Legionella*, suspensions of strains harboring either pSN85 alone (pEV) or pSN85::*legC4* (p*legC4*) induced with IPTG were lysed by boiling in 3X Laemmli buffer. Proteins were separated by SDS-PAGE followed by transfer to polyvinylidene difluoride (PVDF) membrane (ThermoFisher) using a wet transfer cell (BioRad). Membranes were incubated in blocking buffer [5% non-fat milk dissolved in Tris-buffered saline/0.1% Tween 20 (TBST)]. Anti-FLAG (clone M2, Sigma) was diluted at 1:1000 in blocking buffer and incubated with membranes either overnight at 4°C or at ambient temperature for 3 h with rocking. Wash steps were performed 3 times for 10 min each in TBST. HRP-conjugated goat-α-mouse HRP (Sigma) was diluted in blocking buffer at 1:5000 and incubated with membranes for 1-2 hours at room temperature with rocking. Membranes were washed, incubated with ECL substrate (Amersham) and imaged by chemiluminescence using a c300 Azure Biosystems Darkroom Replacer.

#### Statistical analysis

Statistics were performed with GraphPad Prism software using either Mann-Whitney U test or Students’ *t*-test, as indicated, with a 95% confidence interval. In all experiments, error bars denote standard deviation (± S.D.) of samples in triplicates.

## Results

### LegC4 confers a fitness disadvantage on non-flagellated *L. pneumophila* in wild-type mice

We found previously that the *L. pneumophila* effector LegC4 is detrimental to bacterial replication in the mouse lung (30). These experiments were performed using mice and macrophages deficient for production of the NLRC4 inflammasome (*Nlrc4^-/-^*) to prevent flagellin-mediated restriction of *L. pneumophila* replication (10, 30, 42). To confirm that LegC4-mediated phenotypes were not due to loss of NLRC4, we examined fitness of a *legC4*-deficient *L. pneumophila* (Δ*legC4*) strain in the lungs of wild-type C57BL/6 mice using competitive index (CI) experiments. To prevent NLRC4-mediated restriction of bacterial replication we generated flagellin (*flaA*) loss-of-function mutations in our wild-type (Δ*flaA*) and *legC4* mutant (Δ*flaA*Δ*legC4*) strains (see *Materials and Methods*). We also used a previously generated *flaA*::Tn mutant to facilitate selective plating since the transposon confers resistance to chloramphenicol (30). Mice were infected intranasally with a 1:1 mixture of *L. pneumophila* Δ*flaA*Δ*legC4* and *flaA*::Tn for 48 h. Lung tissue was subsequently homogenized and plated on selective media for CFU enumeration and calculation of CI values (see *Materials and Methods*). The Δ*flaA*Δ*legC4* strain significantly outcompeted the *flaA*::Tn mutant in the lungs of wild-type mice, as evidenced by average CI values significantly greater than 1.0 (*P* < 0.01; **Fig 1A**). Our previous study also revealed that expression of *legC4* from a multi-copy plasmid conferred a fitness disadvantage on *L. pneumophila* compared to the wild-type strain (30). To confirm these results in wild-type mice, we generated a strain of *L. pneumophila* Δ*flaA*Δ*legC4* harboring a plasmid encoding *legC4* under control of its endogenous promoter (pJB::p*legC4*). Plasmid expression of *legC4* (pJB::p*legC4*) resulted in significantly impaired fitness defect compared with the Δ*flaA* parental strain, which was not observed for a Δ*flaA*Δ*legC4* strain harboring vector alone (pJB) (*P*<0.05, **Fig 1B**). These data demonstrate that NLRC4 does not affect LegC4-mediated attenuation of *L. pneumophila* replication in the mouse lung. To fully evaluate LegC4-mediated phenotypes, the remainder of our study was performed using *flaA*-deficient strains and bone marrow-derived macrophages (BMDMs) derived from wild-type mice.

**Figure 1.**
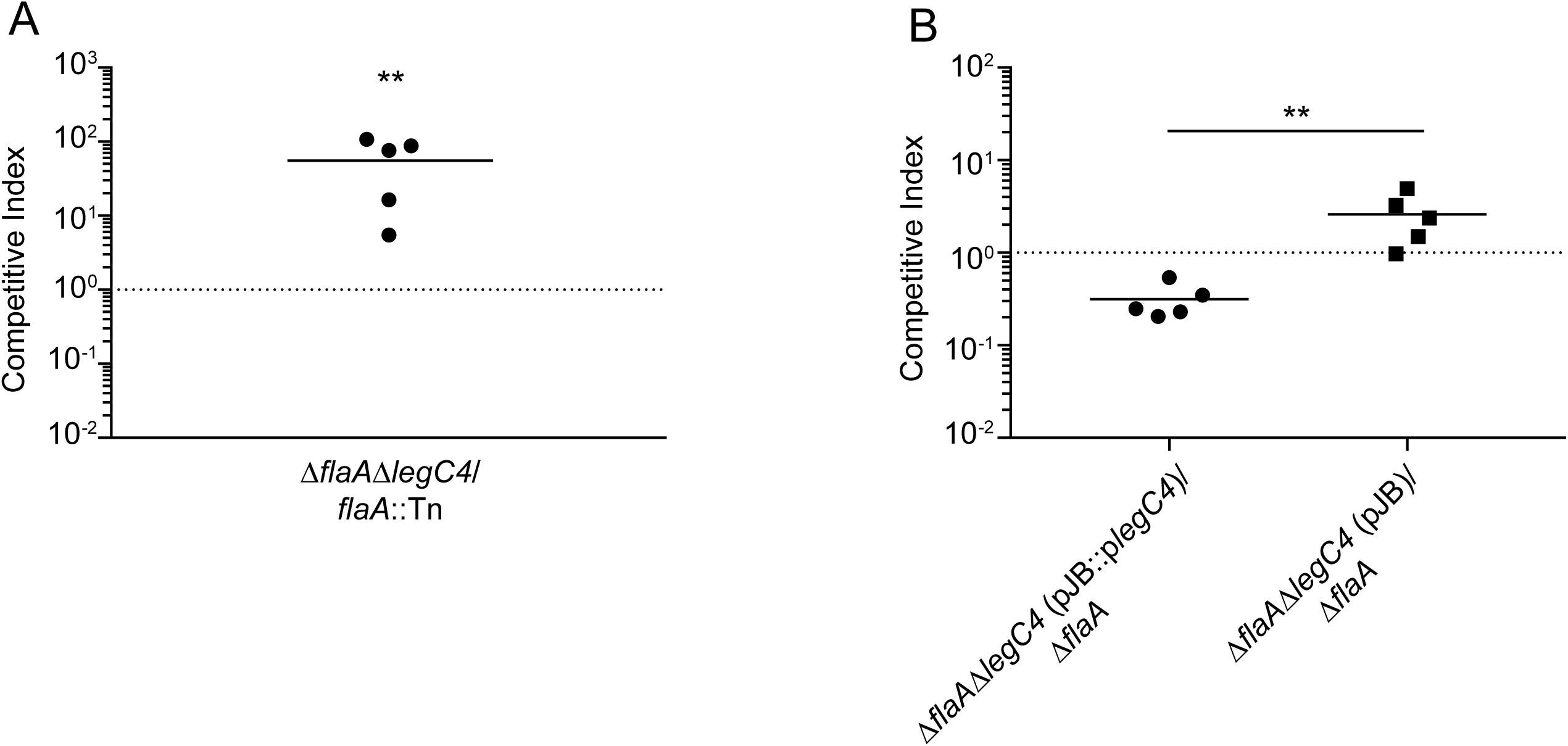
LegC4 attenuates *L. pneumophila* fitness in wild-type mice. (A) Competitive index (CI) *of* Δ*flaA*Δ*legC4* vs. *flaA::*Tn [chloramphenicol(Cm)^R^] from the lungs of wild-type mice. (B) CI of Δ*flaA*Δ*legC4* (pJB::p*legC4*) or *ΔflaA*Δ*legC4* (pJB) vs. Δ*flaA* from the lungs of wild-type mice. Each symbol represents an individual animal and the line represents the mean CI values. Asterisks denote statistical significance by Mann-Whitney U test (\*\**P*<0.01). Data are representative of at least two independent experiments.

### Plasmid expression of *legC4* attenuates *L. pneumophila* replication in BMDMs

Since plasmid expression of *legC4* attenuated *L. pneumophila* fitness in the mouse lung, we examined whether this also occurred in macrophages *ex vivo* using BMDMs derived from wild-type mice. We quantified bacterial replication within BMDMs over 72 h. Consistent with our previous study, loss of endogenous *legC4* (Δ*flaA*Δ*legC4*) does not affect replication of *L. pneumophila* within primary mouse BMDMs compared to the parental strain (Δ*flaA*) (30) (**Fig 2A**). However, *L. pneumophila* Δ*flaA*Δ*legC4* (pJB::p*legC4*) was significantly attenuated for replication in BMDMs at 48 and 72 h p.i. compared to the empty vector control strain (**Fig 2B**). Furthermore, IPTG-induced expression of *legC4* from a plasmid (p*legC4*) also resulted in impaired *L. pneumophila* replication compared to the control strain (pEV) (*P*<0.01, **Fig 2C**). Fitness defects associated with plasmid expression of *legC4* were specific to intracellular replication since replication in rich media *in vitro* was unaffected (data not shown). These data demonstrate that increased levels of LegC4 are detrimental to *L. pneumophila* intracellular replication in macrophages.

**Figure 2.**
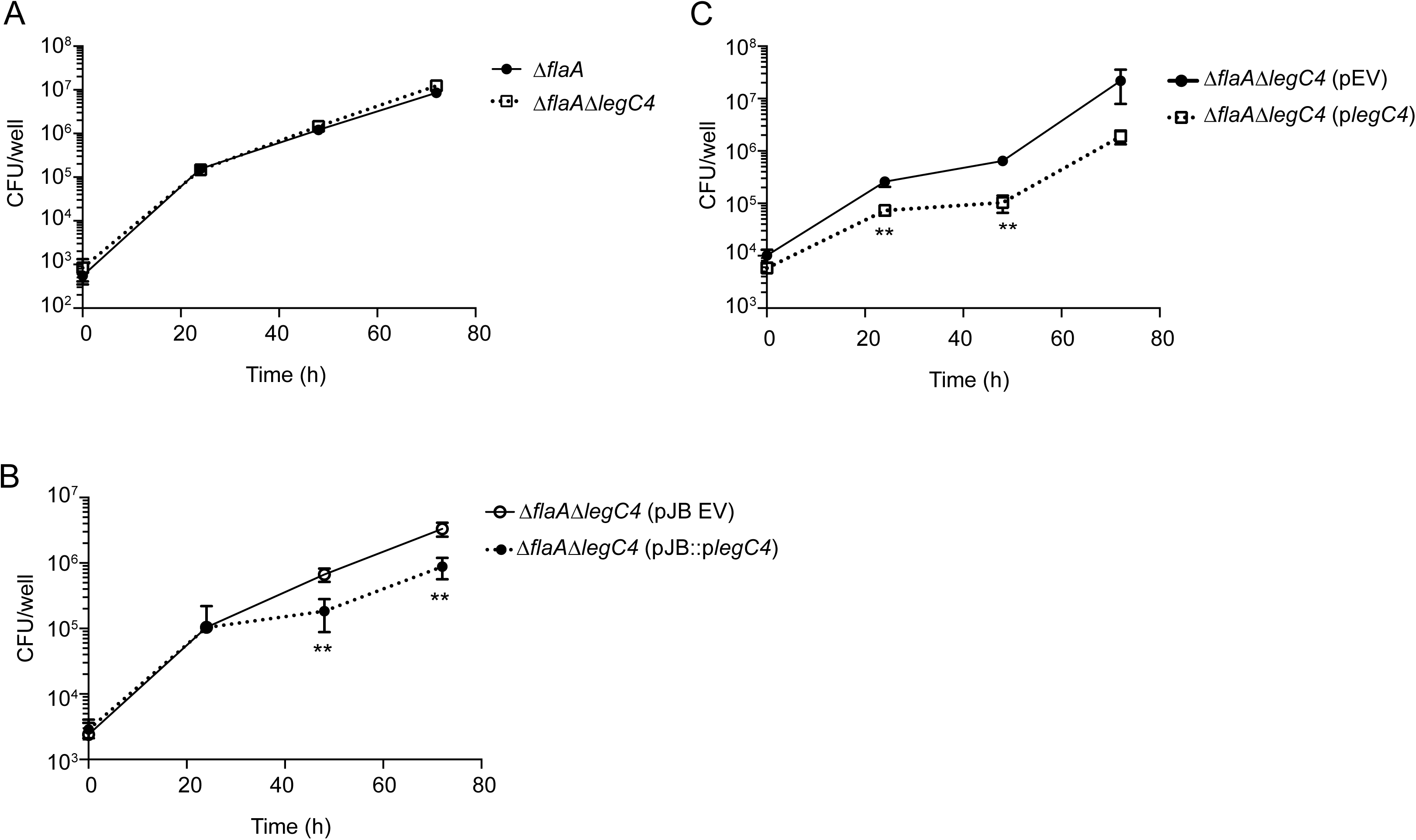
Plasmid expression of *legC4* impairs *L. pneumophila* replication within BMDMs. (A) Growth of *L. pneumophila* **(A)** Δ*flaA* and *ΔflaAΔlegC4*, **(B)** Δ*flaA*Δ*legC4* (pJB) and Δ*flaA*Δ*legC4* (pJB::plegC4) or **(C)** Δ*flaA*Δ*legC4* (pEV) and Δ*flaA*Δ*legC4* (p*legC4*) in BMDMs over 72 h. Expression of *legC4* from p*legC4* was induced with 1 mM IPTG as described (see *Materials & Methods*). Data are shown as mean ± S.D. of samples in triplicates. Asterisks denote statistical significance by Students’ *t*-test (\*\**P*<0.01) and data are representative of three independent experiments.

### LegC4-mediated restriction of *L. pneumophila* replication is dependent on cytokine production

We subsequently investigated the mechanism by which plasmid expression of *legC4* attenuates *L. pneumophila* replication. In BMDMs, *L. pneumophila* infection results in production of pro-inflammatory cytokines through engagement of TLRs by bacterial ligands. Indeed, we previously reported that BMDMs infected with *L. pneumophila* expressing *legC4* secreted increased levels of interleukin (IL)-12 (30). However, increased levels of IL-12 would likely not be sufficient to attenuate *L. pneumophila* intracellular replication within BMDMs. Like IL-12, tumor necrosis factor (TNF) is a pro-inflammatory cytokine expressed downstream of toll-like receptors (TLRs) in macrophages. TNF is important for host defense against *L. pneumophila* in mice and humans and TNF-mediated signaling is sufficient to restrict *L. pneumophila* intracellular replication within macrophages (17-19, 23, 43). Thus, increased TNF signaling could account for LegC4-mediated attenuation of *L. pneumophila* intracellular replication.

We hypothesized that plasmid expression of *legC4* would be sufficient to increase TNF secretion from *L. pneumophila-infected* BMDMs. Wild-type BMDMs were infected with Δ*flaA*, Δ*flaA*Δ*legC4*, Δ*flaA*Δ*legC4* (p*legC4*), or Δ*flaA*Δ*legC4* (pEV) for 6 h and secreted TNF was quantified by ELISA (see *Materials and Methods*). Significantly greater concentrations of TNF were present in the supernatants of cells infected with *L. pneumophila* expressing *legC4* from a plasmid (*P*<0.01, **Fig 3A**). Increased TNF secretion was not due to differences in bacterial replication, since all strains replicated to similar levels at 6 h p.i. (**Fig 3B**). We observed the same phenotype following infection with strains constructed in the Lp02 background, which is a thymidine auxotroph (*thyA^-^*) that is metabolically active but does not replicate in the absence of exogenous thymidine (**Fig S1**). Thus, overexpression of *legC4* results in enhanced TNF secretion from *L. pneumophila*-infected BMDMs.

**Figure 3.**
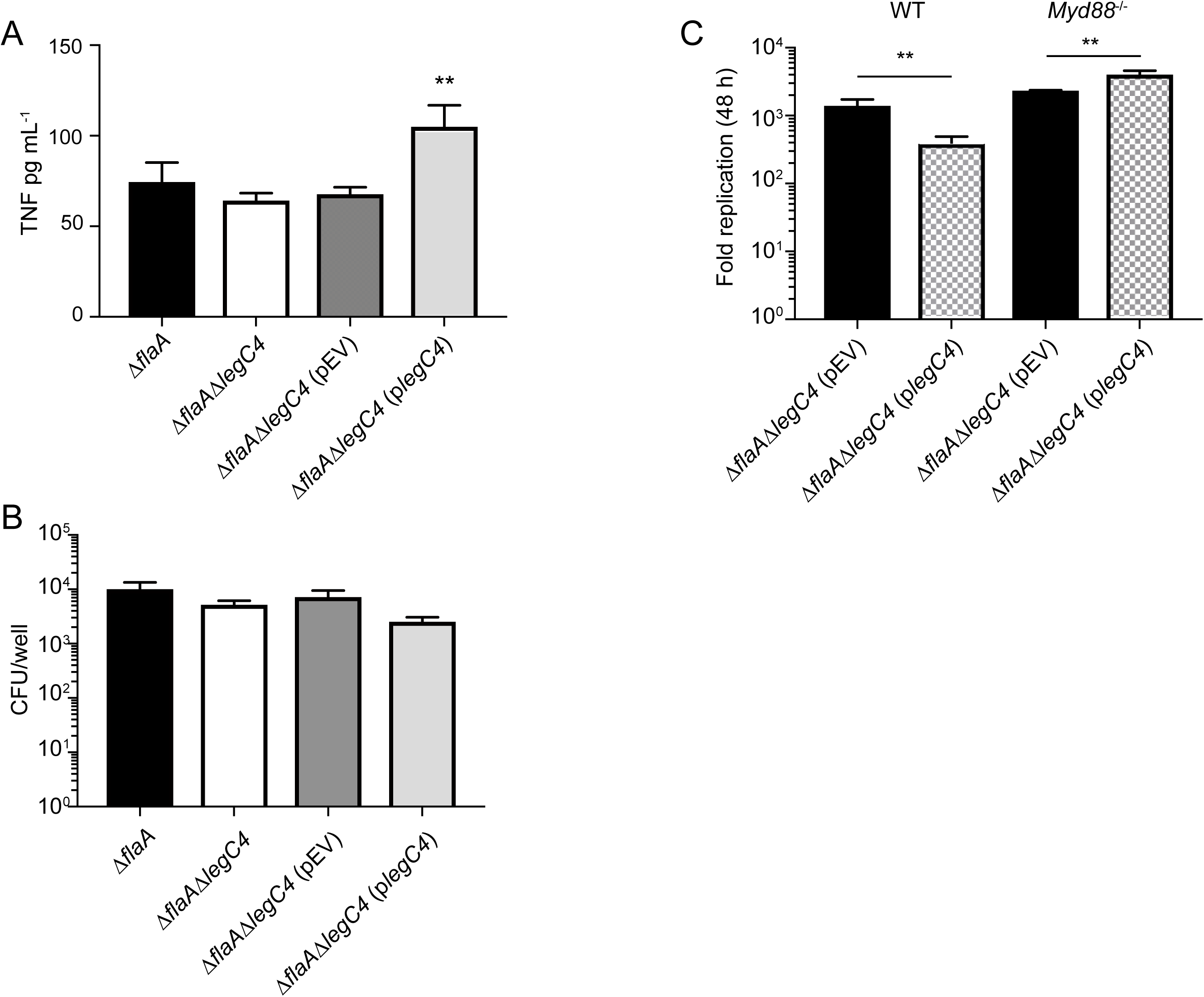
Role of TNF secretion in LegC4-mediated attenuation of *L. pneumophila* replication. **(A)** ELISA for TNF secretion from wild-type BMDMs infected with the indicated strains. (B) Enumeration of *L. pneumophila* strains from BMDMs assayed in (A). **(C)** Fold replication (48 h) of the indicated *L. pneumophila* strains within wild-type or *Myd88^-/-^* BMDMs. Expression of *legC4* was induced with IPTG. Data shown are mean ± S.D. of samples in triplicates. Asterisks denote statistical significance (\*\**P*<0.01) by Students’ *t*-test. Data are representative of at least two independent experiments.

To determine if TNF secretion contributed LegC4-mediated attenuation of intracellular replication, we evaluated *L. pneumophila* replication within *Myd88^-/-^* BMDMs. Since attenuated *L. pneumophila* replication within BMDMs associated with LegC4 were observed at 48 h p.i., we quantified fold replication of the indicated strains at this time point (see *Materials and Methods*). Plasmid expression of *legC4* impaired *L. pneumophila* intracellular replication within wild-type, but not *Myd88^-/-^*, BMDMs (**Fig 3B**). As expected, TNF was not secreted from *Myd88^-/-^* BMDMs under any of our experimental conditions [(44) & data not shown]. Together, these data suggest that pro-inflammatory cytokine production contributes to LegC4-mediated attenuation of *L. pneumophila* intracellular replication in BMDMs.

To further characterize LegC4-mediated restriction of *L. pneumophila* replication within BMDMs, *L. pneumophila* replication was evaluated in the absence of TNF signaling. To determine if TNF signaling contributed to legC4-mediated attenuation of *L. pneumophila* replication, we neutralized TNF in the supernatants of infected wild-type BMDMs using an α-TNF antibody. Wild-type BMDMs were infected with *L. pneumophila* Δ*flaA*Δ*legC4* harboring p*legC4* or pEV in the presence of either α-TNF, Rat IgG isotype control antibody, or neither and fold replication at 48 h p.i. was quantified. Plasmid expression of *legC4* resulted in significantly attenuated *L. pneumophila* replication within untreated and Rat IgG-treated BMDMs (*P*<0.05); however, α-TNF antibody neutralization of TNF restored replication of the *legC4* overexpressing strain to wild-type levels (**Fig 4A**). We subsequently examined replication of these strains in BMDMs deficient for signaling from TNF receptor-1 (TNFR1). *Tnfr1^-/-^* BMDMs were infected with *L. pneumophila* Δ*flaA*Δ*legC4* harboring either p*legC4* or pEV and fold replication was quantified at 48 h p.i. As previously observed, over-expression of *legC4* impaired intracellular replication within wild-type BMDMs (*P*<0.05, **Fig 4B**). However, there was no difference in fold replication of the *legC4*-overexpressing strain compared to the empty vector control within *Tnfr1*^-/-^ BMDMs (**Fig 4B**). Interestingly, overexpression of *legC4* resulted in significantly increased *L. pneumophila* replication within *Tnfr1^-/-^* BMDMs (*P*<0.01, **Fig 4B**). These data demonstrate that TNF signaling contributes to LegC4-mediated attenuation of *L. pneumophila* replication within BMDMs.

**Figure 4.**
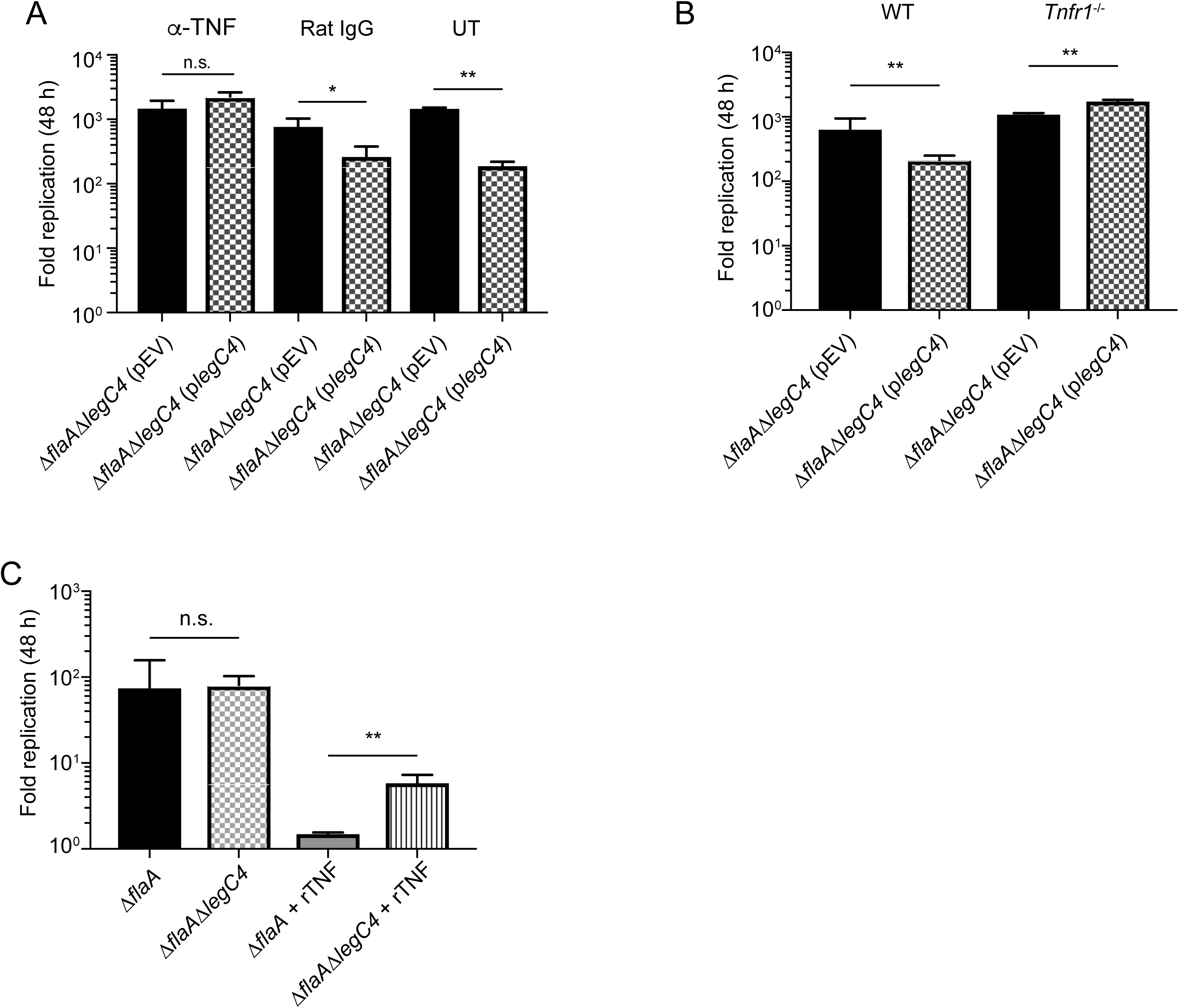
LegC4 augments TNF-mediated restriction of *L. pneumophila* replication. **(A)** Fold replication (48 h) of the indicated *L. pneumophila* strains within wild-type BMDMs treated with 50 ng mL^-1^ α-TNF, isotype control (Rat IgG) or left untreated (see *Materials & Methods*). **(B)** Fold replication (48 h) of the indicated *L. pneumophila* strains within wild-type or *Tnfr1^-/-^* BMDMs. Expression of *legC4* was induced with IPTG. **(C)** Fold replication (48 h) of *L. pneumophila ΔflaA* and Δ*flaA*Δ*legC4* within wild-type BMDMs the presence or absence of 50 ng mL^-1^ recombinant mouse TNF (rTNF). Data shown are mean ± S.D. of samples in triplicates. Asterisks denote statistical significance (*P<0.05, **P<0.01, n.s., not significant) by Students’ *t*-test. Data are representative of at least two independent experiments.

### Endogenous LegC4 exacerbates TNF-mediated restriction of *L. pneumophila* from BMDMs

To further characterize LegC4-mediated restriction of *L. pneumophila* from BMDMs, we examined replication of *L. pneumophila* in BMDMs activated with recombinant mouse TNF (rTNF). Wild-type BMDMs were infected with *L. pneumophila* Δ*flaA* or Δ*flaA*Δ*legC4* in the presence or absence of 50 ng/mL rTNF and fold replication was quantified at 48 h p.i *L. pneumophila* Δ*flaA*Δ*legC4* replicated to significantly greater levels than the parental Δ*flaA* strain in rTNF treated BMDMs (*P*<0.01, **Fig 4C**). As reported above, loss of endogenous *legC4* does not affect *L. pneumophila* replication within untreated BMDMs (**Fig 4C**). These data show that endogenous levels of LegC4 can augment TNF-mediated restriction of *L. pneumophila* replication.

### LegC4 impairs *L. pneumophila* replication in interferon (IFN)-γ activated BMDMs

Interferon (IFN)-γ plays a major role in host defense against *L. pneumophila* in the lung (20, 45). To determine if LegC4-mediated impairment of *L. pneumophila* intracellular replication was specific to TNF, we examined bacterial replication within IFN-γ-activated BMDMs. Fold replication of *L. pneumophila* Δ*flaA*, Δ*flaA*Δ*legC4, ΔflaAΔlegC4* (pEV) or *ΔflaAΔlegC4* (p*legC4*) within wild-type BMDMs activated with recombinant mouse IFN-γ (rIFN-γ) was quantified. We found that overexpression of *legC4* significantly attenuated *L. pneumophila* replication within IFN-γ-activated BMDMs (*P*< 0.05; **Fig 5A**). The *L. pneumophila ΔflaAΔlegC4* mutant replicated to higher levels in IFN-γ-activated BMDMs compared to the parental Δ*flaA* strain (**Fig 5B**). Although the replication difference was not statistically significant (*P*=0.0918), the trend was consistently observed. Together, these data suggest that IFN-γ-mediated restriction of *L. pneumophila* replication is also augmented by LegC4.

**Figure 5.**
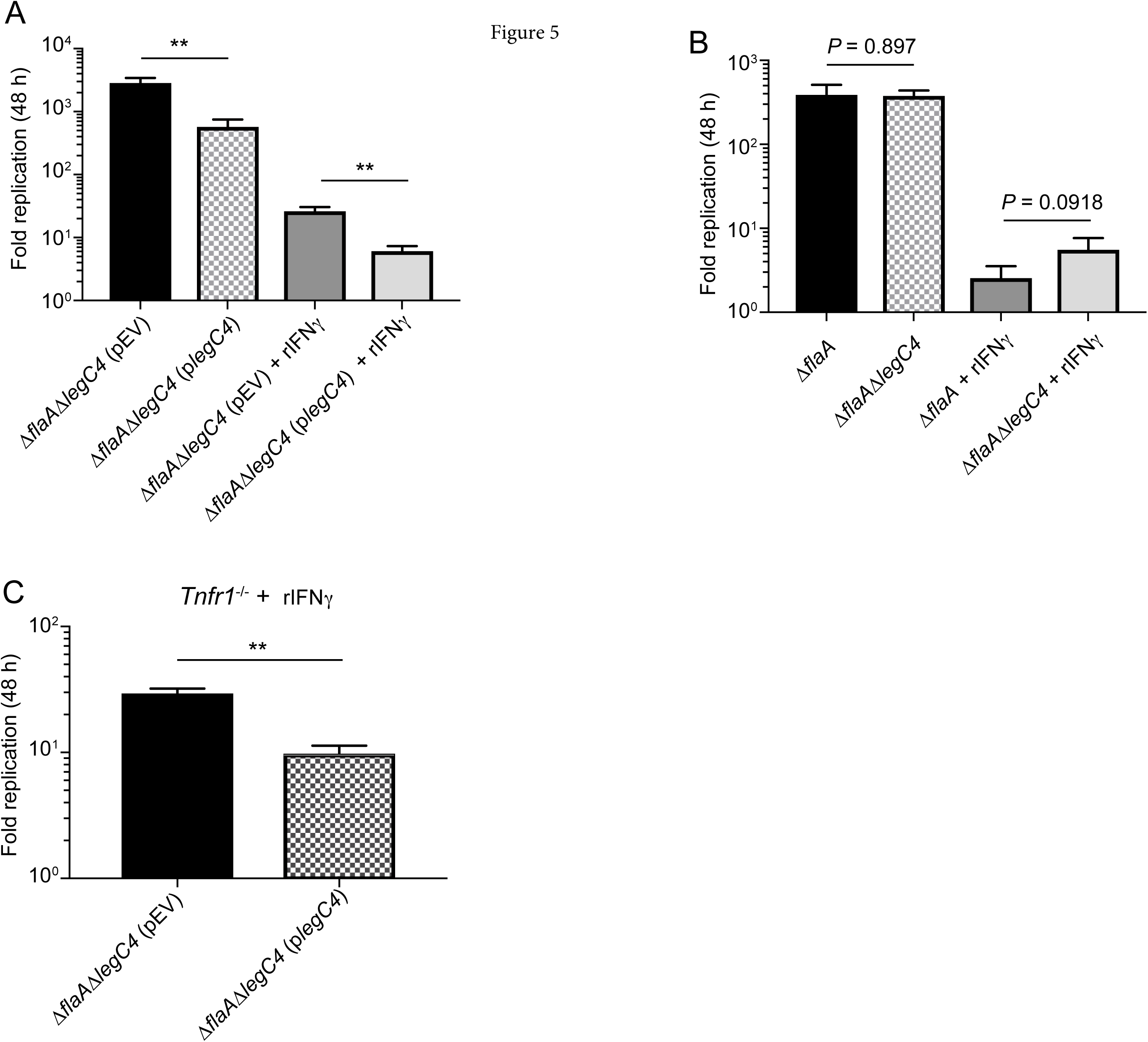
LegC4 enhances IFN-γ-mediated restriction of *L. pneumophila* replication. Fold replication (48 h) of the indicated *L. pneumophila* strains within (**A, B**) wild-type BMDMs or (**C**) *Tnfr1*-/- BMDMs in the presence or absence of 50 ng mL^-1^ recombinant mouse IFN-γ (rIFN-γ) as indicated. Expression of *legC4* was induced with IPTG. Data shown are mean ± S.D. of samples in triplicates. Asterisks denote statistical significance (\*\**P*<0.01) by Students’ *t*-test. Data are representative of at least two independent experiments.

Macrophage activation by IFN-γ results in increased TNF production from macrophages (46, 47). To determine if LegC4-mediated restriction of *L. pneumophila* within IFN-γ activated macrophages was due to TNF signaling, we quantified *L. pneumophila* replication within *Tnfr1^-/-^* BMDMs treated with IFN-γ. Overproduction of LegC4 resulted in significantly decreased *L. pneumophila* replication in IFN-γ-activated TNFR1^-/-^ BMDMs compared to the control strain (*P*<0.001; **Fig 5C**). Thus, LegC4 can also augment non-TNF-mediated restriction of *L. pneumophila* replication through IFN-γ.

### LegC4 impairs *L. longbeachae* replication within cytokine-activated BMDMs

*Legionella longbeachae* replicates within eukaryotic phagocytes using a Dot/Icm secretion system (39). Although the *L. longbeachae* Dot/Icm secretion system is highly similar to that of *L. pneumophila*, their effector repertoires are quite distinct and *L. longbeachae* does not encode a homolog of *legC4* (48, 49). Importantly, *L. longbeachae* is more virulent than *L. pneumophila* and lethal in a mouse model of infection. To determine if LegC4 could attenuate bacterial replication in non-*pneumophila Legionella*, we generated *L. longbeachae* strains either expressing *legC4* (*plegC4*) or harboring the empty vector (pEV). Wild-type BMDMs were infected with these *L. longbeachae* strains and fold-replication at 48 h was quantified. Expression of *legC4* did not impair *L. longbeachae* replication within BMDMs; however, *legC4* expression did result in significantly attenuated *L. longbeachae* replication within rTNF- and rIFN-γ-treated BMDMs compared to the control strain (*P*<0.001; **Fig 6**). These data suggest that LegC4 can augment cytokine-mediated restriction of non-*pneumophila Legionella* within BMDMs.

**Figure 6.**
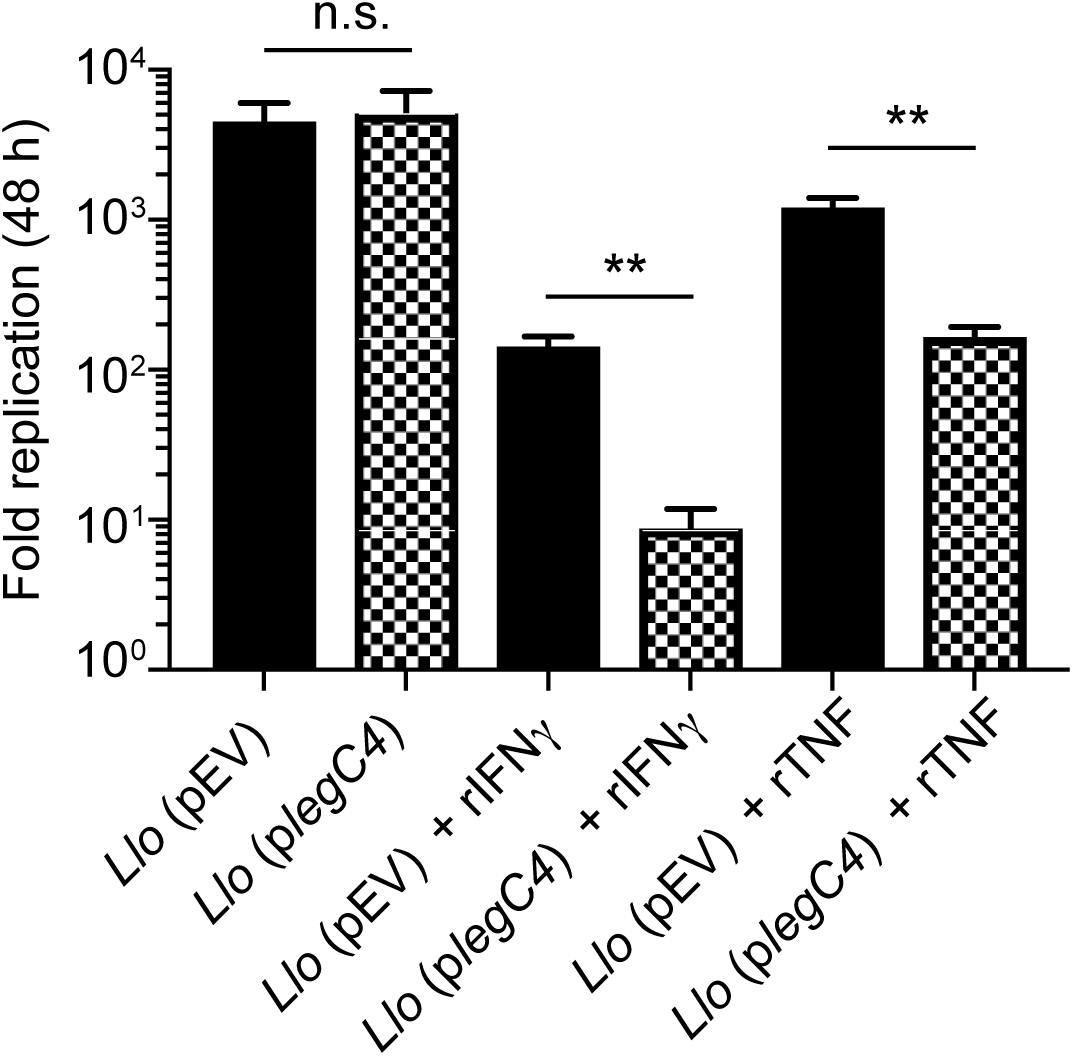
Replication of *L. longbeachae* producing LegC4 in cytokine-treated BMDMs. Fold replication (48 h) of the *L. longbeachae* (*Llo*) harboring the indicated plasmids within wild-type BMDMs the presence or absence of 50 ng mL^-1^ rTNF or rIFN-γ, as indicated. Expression of *legC4* was induced with IPTG. Expression of *legC4* was induced with IPTG. Data shown are mean ± S.D. of samples in triplicates. Asterisks denote statistical significance (\*\**P*<0.01, n.s., not significant) by Students’ *t*-test. Data are representative of at least two independent experiments.

## Discussion

The data presented in this study support the hypothesis that LegC4 potentiates cytokine-mediated host defense against *Legionella*. Our previous work (30) identifying LegC4 as contributing to *L. pneumophila* clearance from the lung was performed using flagellated *L. pneumophila* in a NLRC4-deficient (*Nlrc4^-/-^*) mouse model. To fully evaluate the mechanisms of LegC4-mediated clearance, utilized wild-type mice and BMDMs. Consistent with our previous study (30), we found that loss-of-function mutation in the *legC4* gene (Δ*legC4*) conferred a fitness advantage on *L. pneumophila* Δ*flaA* within the wild-type mouse lung. Moreover, complementation of the Δ*legC4* mutation by a plasmid encoding *legC4 in trans* conferred a fitness disadvantage on *L. pneumophila* compared to the parental strain. Also consistent with our previous report, *L. pneumophila* Δ*flaA*Δ*legC4* replication within BMDMs did not differ from replication of the Δ*flaA* strain. However, in the present study, we found that plasmid expression of *legC4* was sufficient to attenuate *L. pneumophila* replication within BMDMs. Although *legC4* was expressed downstream of its endogenous promoter, an exaggerated phenotype likely occurred due to expression from a multi-copy plasmid, suggesting a potential dose-response. Importantly, this strain provided us with a tool to increase the magnitude of LegC4-mediated fitness attenuation within cultured cells. These phenotypes were corroborated by the observation that endogenous LegC4 was deleterious in cytokine-activated BMDMs. We further found that the fitness disadvantage associated with plasmid expression of *legC4* was abolished in macrophages deficient for TNF-mediated signaling, suggesting that LegC4 is able to exacerbate cytokine-mediated antimicrobial responses. Finally, we determined that LegC4 could impair replication of *L. longbeachae* in cytokine-activated macrophages. Together, these data suggest that LegC4 potentiates cytokine-mediated restriction of *L. pneumophila* within macrophages.

Inflammation is mediated primarily through cytokine secretion, which is critical for restriction of *L. pneumophila* replication in vivo. This has been evidenced by the inability of *Myd88^-/-^* mice to control *L. pneumophila* replication. Specifically, *Myd88^-/-^* BMDMs will not secrete TNF during infection. The inability of plasmid-expressed *legC4* to attenuate *L. pneumophila* replication in *Myd88^-/-^* BMDMs is likely due to lack of TNF signaling. In addition, LegC4-mediated increases in TNF secretion may amplify *Tnf* expression, which would further restrict *L. pneumophila* replication.

Pro-inflammatory cytokines contribute to host defense against *L. pneumophila in vivo* and in cultured macrophages (8, 50). Mice deficient for TNF-mediated signaling have increased pulmonary bacterial burdens and can succumb to infection (23, 43). TNF can signal through both TNFR1 and TNFR2; however, TNFR1-mediated signaling is primarily responsible for *L. pneumophila* restriction within alveolar macrophages *in vivo* (23) and is potentiated by LegC4. In the lung, multiple cell types contribute to TNF production, a consequence of which would be higher local TNF concentrations (16, 18, 23, 51). In addition, production of IFN-γ during *L. pneumophila* infection *in vivo* is mediated primarily by circulating natural killer (NK) cells (45, 52).

Our observation that the *L. pneumophila* Δ*legC4* mutant had fitness advantage compared to wild-type in the mouse lung but not in cultured macrophages suggested that LegC4 was detrimental to replication under specific environmental conditions. This was supported by the observation that attenuated *L. pneumophila* replication was correlated with increased TNF secretion from BMDMs. Since the *L. pneumophila-infected* lung is an inflammatory environment, we examined whether cytokine-mediated restriction was exacerbated by LegC4. Abrogation of signaling from TNFR1 was sufficient to alleviate LegC4-mediated restriction of intracellular replication. Increased replication of the Δ*legC4* mutant within rTNF-treated BMDMs strongly suggests that pro-inflammatory responses are exacerbated by LegC4. This conclusion was corroborated by the observation that the Δ*legC4* mutant consistently replicated to higher levels within IFN-γ-activated macrophages compared to untreated macrophages.

Similar to *L. pneumophila, L. longbeachae* replicates within an LCV by employing a Dot/Icm secretion system and a repertoire of translocated effector proteins (39). Despite high levels of homology between the Dot/Icm secretion systems of these two organisms, the effector repertoires are quite diverse and *L. longbeachae* does not encode a homolog of *legC4* (48, 49). In contrast to *L. pneumophila, L. longbeachae* is highly virulent in a mouse model of Legionnaires’ disease (53, 54). Lethality in mice is likely due to *L. longbeachae* being poorly immunostimulatory and failing to induce substantial levels pro-inflammatory cytokines during infection. However, pro-inflammatory cytokines contribute to host defense against *L. longbeachae* in BMDMs and in vivo (53). Since inter-species translocation of Dot/Icm effectors by *Legionella* has been previously observed (39, 55), we introduced *legC4* into *L. longbeachae*. Production of LegC4 by *L. longbeachae* resulted in significantly attenuated replication within cytokine treated, but not untreated, BMDMs. These data reinforce our previous observations and demonstrate that LegC4-mediated restriction is not specific to *L. pneumophila*. Since *L. longbeachae* infection does not induce appreciable TNF secretion from BMDMs (53), it is likely that the concentration of TNF secreted by these cells is too low to permit LegC4-mediated restriction. Together with relatively low levels of effector translocation by *L. longbeachae* compared to *L. pneumophila* (39), the amount of translocated LegC4 may be insufficient to restrict bacterial replication within untreated BMDMs. However, LegC4 is sufficient to attenuate *L. longbeachae* replication within BMDMs activated with either rTNF or rIFN-γ. Whether LegC4 can protect mice from *L. longbeachae*-mediated lethality will be the subject of a future study.

Multiple effectors contribute to the innate immune response to *L. pneumophila* infection [reviewed in (56)]. Together with our data, these studies point to a complex interplay between effectors during *Legionella* infection of mammalian host. The effectors LnaB and LegK1 enhance NF-κB activation, which augments immune signaling (57, 58). Since mammals are a dead-end host for *Legionella*, the evolutionary basis for effector modulation of NF-κB is intriguing. Interestingly, the effector EnhC enhances *L. pneumophila* replication in TNF-activated macrophages (59), the opposite of what we have observed for LegC4. Thus, it is tempting to speculate that there may be interplay between EnhC and LegC4 within *L. pneumophila* infected cells. Future investigations will reveal whether LegC4-mediated phenotypes are dependent on other Dot/Icm-translocated effectors.

In summary, we found that the Dot/Icm effector LegC4 can augment cytokine-mediated restriction of *Legionella* replication within macrophages. These data add to the growing body of literature on effector triggered immunity in animal cells. As an accidental pathogen that did not co-evolve under the selective pressure of an innate immune system, *L. pneumophila* continues to provide insight into novel mechanisms of innate immunity towards intracellular bacterial pathogens. Consequently, further understanding of LegC4 function will reveal strategies to augment pro-inflammatory signaling. Thus, this study has provided the foundation for future investigations into the molecular mechanism by which LegC4 enhances host defense against intracellular bacterial pathogens.

## Acknowledgements

We would like to thank Drs. B.A. Montelone and A. L. Passarelli for critical reading of the manuscript. The idea for a portion of this work was conceived in Dr. Craig Roy’s laboratory at the Yale School of Medicine. This work was supported by start-up funds and a Mentoring Award from Kansas State University (to S.R.S.), an NIH K-INBRE Developmental Research Project Award (P20GM103418; to S.R.S), and an NIH K-INBRE Semester Scholar Award (to A.J.H.). The funders had no role in study design, data collection and interpretation, or the decision to submit the work for publication.

## Supplemental information

**Figure S1.**
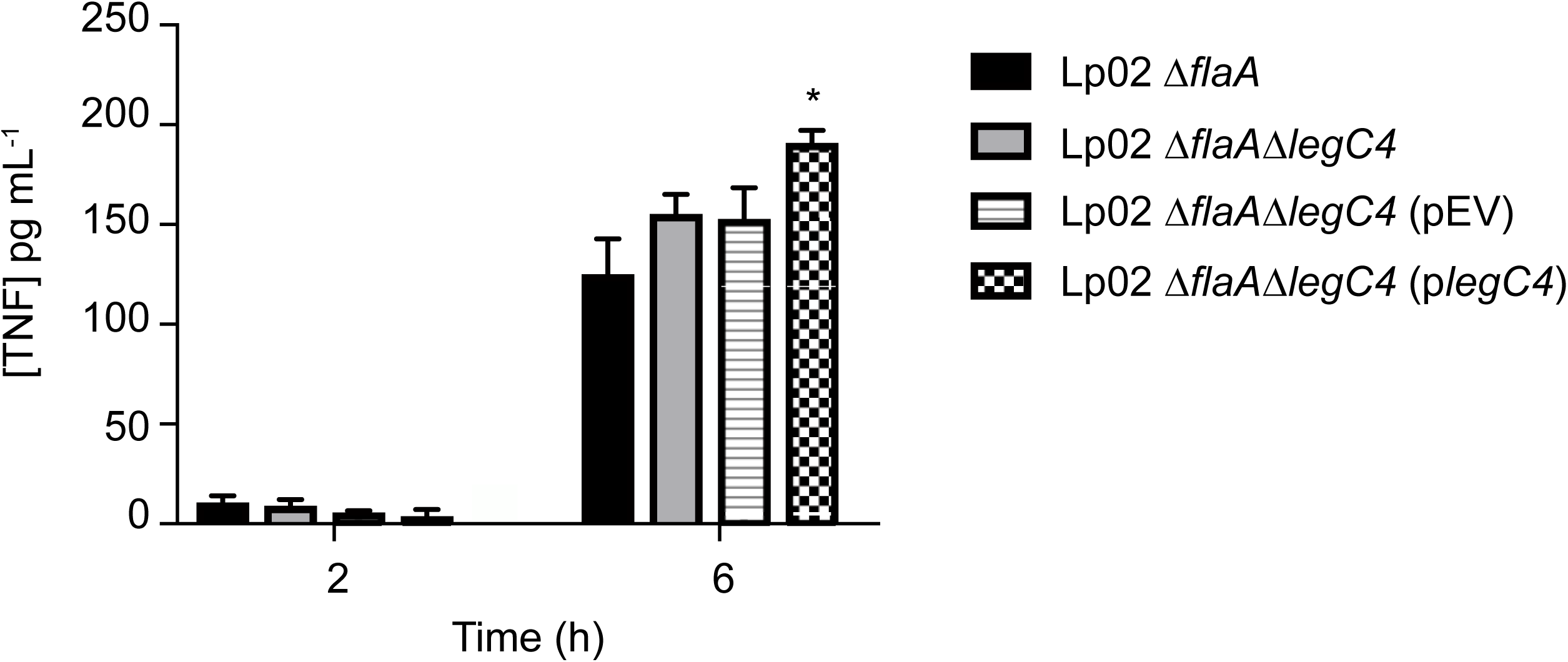
TNF secretion from BMDMs infected with Lp02 strains. (A) ELISA for TNF secreted from wild-type BMDMs infected the indicated Lp02 strains for 2 h or 6 h in the absence of exogenous thymidine. Asterisks denote statistical significance (\**P*<0.05) by Students’ *t*-test. Data are representative of two independent experiments.

